# Molecular basis of CTCF binding polarity in genome folding

**DOI:** 10.1101/2019.12.13.876177

**Authors:** Elphège P. Nora, Laura Caccianini, Geoffrey Fudenberg, Vasumathi Kameswaran, Abigail Nagle, Alec Uebersohn, Kevin So, Bassam Hajj, Agnès Le Saux, Antoine Coulon, Leonid A. Mirny, Katherine S. Pollard, Maxime Dahan, Benoit G. Bruneau

## Abstract

Current models propose that boundaries of mammalian topologically associating domains (TADs) arise from the ability of the CTCF protein to stop extrusion of chromatin loops by cohesin proteins (Merkenschlager & Nora, 2016; Fudenberg, Abdennur, Imakaev, Goloborodko, & Mirny, 2017). While the orientation of CTCF motifs determines which pairs of CTCF sites preferentially stabilize DNA loops (de Wit et al., 2015; Guo et al., 2015; Rao et al., 2014; Vietri Rudan et al., 2015), the molecular basis of this polarity remains mysterious. Here we report that CTCF positions cohesin but does not control its overall binding or dynamics on chromatin by single molecule live imaging. Using an inducible complementation system, we found that CTCF mutants lacking the N-terminus cannot insulate TADs properly, despite normal binding. Cohesin remained at CTCF sites in this mutant, albeit with reduced enrichment. Given that the orientation of the CTCF motif presents the CTCF N-terminus towards cohesin as it translocates from the interior of TADs, these observations provide a molecular explanation for how the polarity of CTCF binding sites determines the genomic distribution of chromatin loops.

## Main text

In mammalian genomes, cohesin complexes accumulate at CTCF binding sites, and cohesin-dependent chromatin loops preferentially engage pairs of CTCF sites with convergent motif orientation (Rao et al., 2014; Vietri Rudan et al., 2015). Inverting one CTCF motif can lead to repositioning of the corresponding DNA loop (Bemmel et al., 2019; de Wit et al., 2015; Guo et al., 2015; Sanborn et al., 2015). Two non-exclusive models may account for both localization of cohesin at CTCF sites and directional DNA looping. Cohesin could load at CTCF binding sites, downstream of the motif, and initiate loop extrusion unidirectionally (Nichols & Corces, 2015). Alternatively, cohesin could load throughout TADs and translocate bi-directionally as it extrudes DNA loops, only stopping when it encounters a CTCF sites in the proper orientation (Fudenberg et al., 2016; Sanborn et al., 2015).

To test these models, we first measured the impact of depleting CTCF on cohesin binding and positioning on chromosomes. As previous studies using inducible CTCF knock-out reported that cohesin can still bind 80% of initial sites even after 10 days (Busslinger et al., 2017), we sought to achieve more efficient depletion. Using a mouse embryonic stem cell (mESC) line in which CTCF can be degraded by the auxin inducible degron (AID) system (Nora et al., 2017), we observed near-complete disappearance of the cohesin ring subunit RAD21 by ChIP-seq from its initial position at CTCF peaks (**Figure 1a, b**). However, spike-in calibration revealed identical amount of chromatin pulled down by a RAD21 antibody in the absence of CTCF (**Figure 1c**). Thus, while cohesin no longer accumulates at CTCF sites in the absence of CTCF, it still associates with chromatin, indicating it must be redistributed away from CTCF sites – supporting the translocation-and-block model.

**Figure 1.**
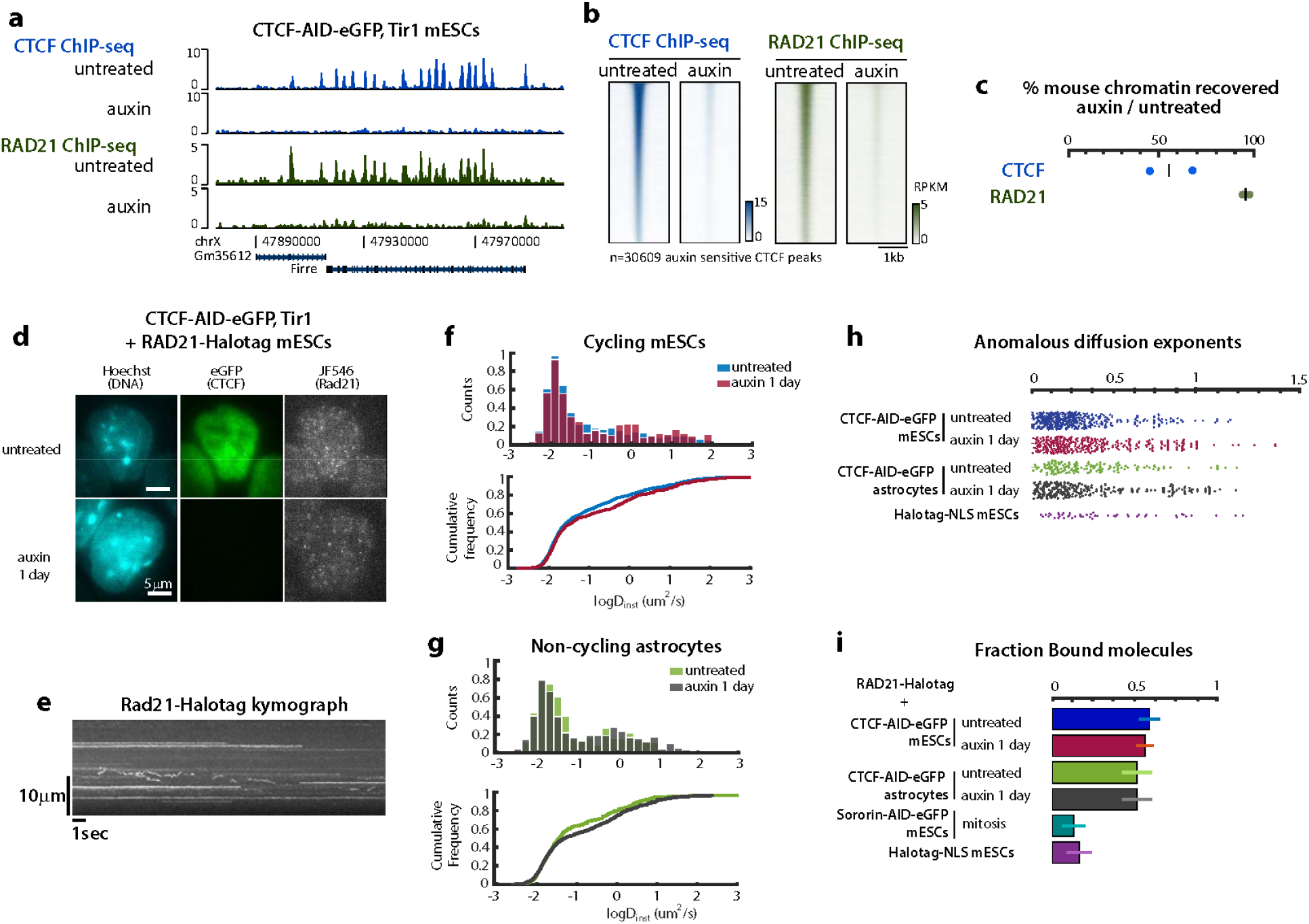
CTCF acts as a positioning but not a loading factor for cohesin. **a** and **b**, Rad21 ChIP-seq enrichment at CTCF peaks is lost after depleting CTCF in CTCF-AID mESCs. **c**, Percentage of ChIP-seq reads mapping to mouse versus spike-in (Drosophila) genomes, using antibodies against either mouse CTCF or mouse Rad21 in CTCF-AID mESCs, normalized to values obtained before CTCF depletion by auxin. Each replicate is plotted separately. **d**. HILO imaging of single endogenous cohesin molecules in live CTCF-AID RAD21-Halotag knockin mESCs labeled with limiting JF546 ligand (50ms acquisitions). **e**, Part of a kymograph generated by an xy line scan across a single cell, illustrating the various diffusion behaviors of RAD21-Halotag in mESCs (50ms acquisitions). **f**, The distribution of diffusion coefficients (D_inst_) of Rad21 molecules, although significantly different statistically, is only mildly altered by CTCF depletion. This slight increase in the number of fast diffusing molecules is observed both in cycling mESCs and **g**, non-cycling astrocytes. 50ms acquisitions. KS test: p=0.0196 for mESCs and p=0.0014 for astrocytes, pooling trajectories from all cells. **h**, anomalous diffusion exponents of RAD21 trajectories (50ms acquisitions) indicating that Rad21 molecules imaged are overwhelmingly sub-diffusive (<1). **i**, CTCF depletion does not alter the fraction of bound cohesin (5ms acquisitions). Mitotic block triggered by depleting Sororin serves as a positive control for free diffusing cohesin, since most cohesin is unloaded in prophase. Means with standard deviation. See Methods for detailed statistics.

To directly visualize how loss of CTCF may affect cohesin dynamics and association with DNA, we performed single molecule tracking of RAD21 in WT (**Figure 1 – figure supplement 1a-g**) and CTCF-AID mESCs (**Figure 1d-g**) by targeting both *Rad21* alleles with a HaloTag. Consistent with previous measurements (Hansen, Pustova, Cattoglio, Tjian, & Darzacq, 2017), 60% of RAD21 molecules were bound to chromatin (**Figure 1i**). Depleting CTCF did not affect this fraction, nor the distribution of diffusion coefficients or the anomalous diffusion exponent of RAD21 (**Figure 1f-i**). Cell cycle and sister chromatid cohesion were not a confounding effect in these imaging modalities (see Methods) since we obtained similar results in each single cycling mESCs (**Figure 1 – figure supplement 1g**), and in non-cycling astrocytes (**Figure 1f-g**). However, CTCF depletion led to a modest but reproducible increase in the number of fast-diffusing molecules (−1 < LogD_inst_ < 0), in both cycling and non-cycling cells (**Figure 1f-g**). These fast diffusing molecules were nevertheless not completely free, since they diffused more slowly than unbound cohesin (LogDinst > 0), as estimated from imaging cells blocked in early M phase by means of a 6-hr depletion of Sororin (**Figure 1 – figure supplement 1h-o**). Taken together with the spike-in ChIP-seq, our results refute the idea that CTCF promotes bulk loading of cohesin and support a mechanism whereby CTCF acts by blocking translocating cohesin.

We investigated how CTCF blocks cohesin translocation and triggers TAD insulation. Mutational analysis of CTCF is challenging because CTCF is essential for long-term cell survival (Nora et al., 2017; Sleutels et al., 2012), and mutations altering CTCF protein stability or CTCF binding will *de facto* alter cohesin positioning and TAD folding – since insulation of TADs relates quantitatively to CTCF levels (Nora et al., 2017). To overcome these obstacles, we used a complementation system where inducible CTCF cDNA transgenes are stably targeted in CTCF-AID cells, so that auxin degrades endogenous CTCF and doxycycline triggers expression of the CTCF transgene (**Figure 2a**). Precise comparison of expression levels between cell lines was achieved by flow-cytometry for mRuby2, fused in frame to transgenic CTCF. TAD folding was surveyed across all genotypes by Chromosome Conformation Capture Carbon-Copy (5C) using a previously validated design (Nora et al., 2017). To calibrate our assay, we analyzed two independent lines expressing the full-length CTCF cDNA at either high or low level, together with one cell line not expressing the transgene. Insulation (**Methods**) scaled linearly with transgene expression (Figure 2b, dashed line). Expression of the full-length transgene (high) was approximately one fifth of endogenous CTCF-AID-eGFP, which is less than half untagged CTCF (**Figure 2 – figure supplement 2a-c**).

**Figure 2.**
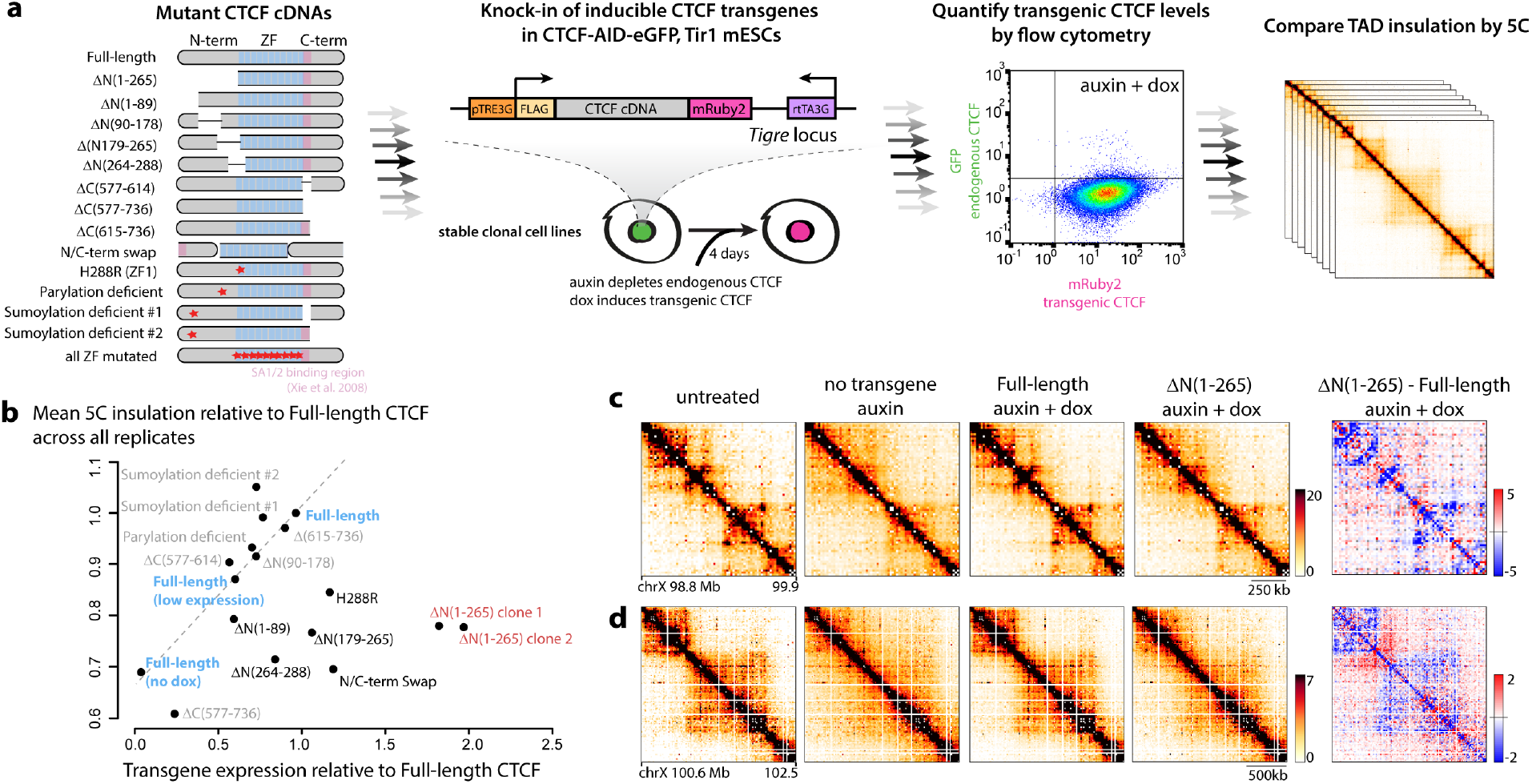
Deletion scanning reveals aminoterminal domains of CTCF mediate TAD folding. **a**, Experimental pipeline for the mutational analysis of CTCF using stable transgenic mESCs. Red stars indicate amino-acid substitutions as detailed in the Methods. Flow cytometry confirming homogenous induction of the transgenes (only full-length is shown). Gates were placed using untagged cells. **b**, Summary of 44 5C experiments across 16 stable mESC lines treated with dox and auxin. Each data point is the mean of the insulation scores at the 6 TAD boundaries initially detected in WT untreated samples, averaged across at least two 5C replicates, and presented as ratios relative to insulation measured in the full-length CTCF transgene. Transgene expression values correspond to flow-cytometry means across at least two replicates. The dashed line is the linear regression indicating how insulation depends on transgene expression for full-length CTCF, and was obtained by comparing cell lines with either high, low or no full-length transgene expression. Transgenes with all zinc fingers (ZFs) mutated were poorly expressed and not assessed by 5C. **c**, and **d** snapshots of 5C data binned at 15kb, and corresponding differential heatmaps highlighting folding defects in the Δ(1-265) mutants.

We first deleted C(577-614), which contains a region expected to mediate the interaction between CTCF and cohesin based on *in vitro* data (Xiao, Wallace, & Felsenfeld, 2011), and encompasses the C-terminal internal RNA-binding region, RBRi (**Figure 2 – figure supplement 1d and 2a**) (Hansen et al., 2018; Saldaña-Meyer et al., 2014). ΔC(577-614) is expressed at around 60% of level of the full length transgene, confirming the region contributes to CTCF stability (**Figure 2 – figure supplement 2b**)(Hansen et al., 2018). ΔC(577-614) displayed lower DNA binding by ChIP-seq (**Figure 2 – figure supplement 2e-g**) and rescued insulation as expected based on its expression level (**Figure 2b and Figure 2 – figure supplement 2c-d**). Altogether, C(577-614) appears dispensable for functionally connecting CTCF and cohesin *in vivo*, and contributes minimally to TAD folding beyond promoting CTCF binding, at least in the region surveyed. Another CTCF domain must therefore mediate cohesin blocking and directional loop retention.

We proceeded to establish an additional 12 stable cell lines, each harboring a different mutated CTCF cDNA, leaving the core of the DNA binding domain intact (central Zinc-finger array, or ZF - **Figure 2a and Figure 2 – figure supplement 1d**). Several CTCF mutants failed to rescue TAD insulation to the extent expected from their expression levels (**Figure 2b**). Deletion of the entire N-terminal domain ΔN(1-265) had the most dramatic impact (**Figure 2c-d**). Within the N-terminus multiple sub-regions participate to the ability of CTCF to insulate TADs (**Figure 2b**): ΔN(1-89) triggered a mild but detectable insulation defect, while ΔN(179-265) had a more pronounced effect. ΔN(264-288), which overlaps one RNA-binding region and ZF1, as well as mutation of the ZF1 itself (H288R), also led to insulation defects and is characterized further in a parallel study (Saldana-Meyer et al., 2019).

To understand the chromatin folding defects in ΔN(1-265) we measured binding of transgenic CTCF and endogenous Rad21 by ChIP-seq. Deleting the entire N-terminus did not alter CTCF binding, as indicated by FLAG pull-down (**Figure 3**). RAD21 enrichment at FLAG-CTCF peaks remained detectable in the ΔN(1-265) mutant, but was reduced two-fold (**Figure 3**). Therefore, proper retention of cohesin at CTCF sites requires N(1-265), indicating that the CTCF N-terminus either participates in inhibiting cohesin translocation (thereby promoting insulation), or – non-exclusively – protects blocked cohesin from unloading (thereby bolstering 5C peaks between CTCF sites - **Figure 4 – figure supplement 1**).

**Figure 3.**
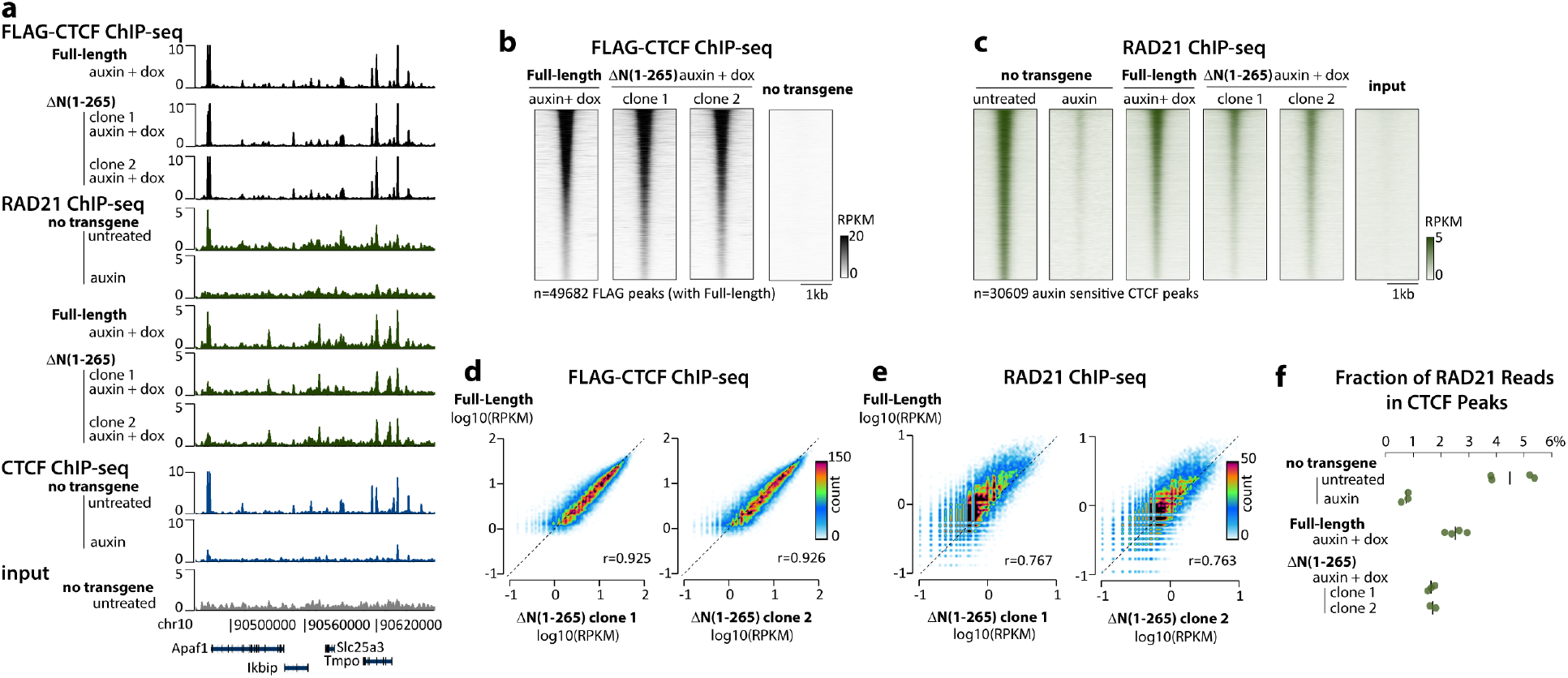
The CTCF N-terminus participates in but is not strictly required for cohesin positioning at CTCF sites. **a**, ChIP-seq tracks snapshot, **b-c**, density plots **d-e**, scatter plots and **f**, Fraction of Reads in Peak (FRIP) scores indicates that RAD21 is still detected at CTCF peaks(Nora et al., 2017) in cells expressing CTCF ΔN(1-265), albeit with a two-fold reduced enrichment compared to the full-length CTCF transgene.

Given that deleting the CTCF N-terminus led to milder insulation defects than complete CTCF depletion, and that deleting the C-terminus had little to no effect, the Zinc-finger array mediates some degree of insulation and must therefore participate in halting cohesin translocation. The Zinc finger domain confers to CTCF an unusually long residence time to CTCF for a transcription factor (Agarwal, Reisser, Wortmann, & Gebhardt, 2017; Hansen et al., 2017), as well as uniquely distorts DNA (MacPherson & Sadowski, 2010) and positions nucleosomes (Fu, Sinha, Peterson, & Weng, 2008) in a fashion that might interfere with loop extrusion by cohesin.

The importance of the CTCF N-terminus draws support from evolutionary data: while the ZF domain of CTCF is highly conserved across bilateria (Heger, Marin, & Schierenberg, 2009), vertebrate and invertebrate N-termini are highly divergent. In *Drosophila*, CTCF binding sites also overlap cohesin ChIP-seq peaks (Li et al., 2015) (**Figure 3 – figure supplement 1**), but do not exhibit motif orientation bias at domain borders (Matthews & White, 2019) and do not anchor Hi-C peaks (Cubeñas-Potts et al., 2016; Eagen, Aiden, & Kornberg, 2017). This reinforces further the notion that, while the conserved ZF domain is an impediment to cohesin translocation, the mammalian N-terminus is required to fully retain cohesin and stabilize chromatin loops as they appear by Hi-C. While the CTCF N-terminus is highly conserved across mammals, it is highly divergent from that of its paralog BORIS/CTCFL, which is therefore not anticipated to share the functions of CTCF in genome architecture (Debruyne et al., 2019; Pugacheva et al., 2015).

Altogether, our observations also explain why TAD boundaries are preferentially populated by pairs of CTCF sites with binding sites in a convergent orientation, and why inverting a CTCF site impairs chromatin interactions, in spite of leaving cohesin ChIP enrichment unchanged (de Wit et al., 2015). Indeed, orientation of the CTCF motif ensures that cohesin translocating from the inner portion of TADs encounters the N-terminus of CTCF (**Figure 4g and Figure 4 – figure supplement 1**). When the N-terminus is placed C-terminally of the Zinc finger array, CTCF is unable to rescue TAD folding, indicating that oriented presentation of the N-terminus is crucial (**Figure 2b**). These results pave the way for further mechanistic dissection of the process, and will guide investigation of the molecular details of how CTCF and cohesin interact.

**Figure 4.**
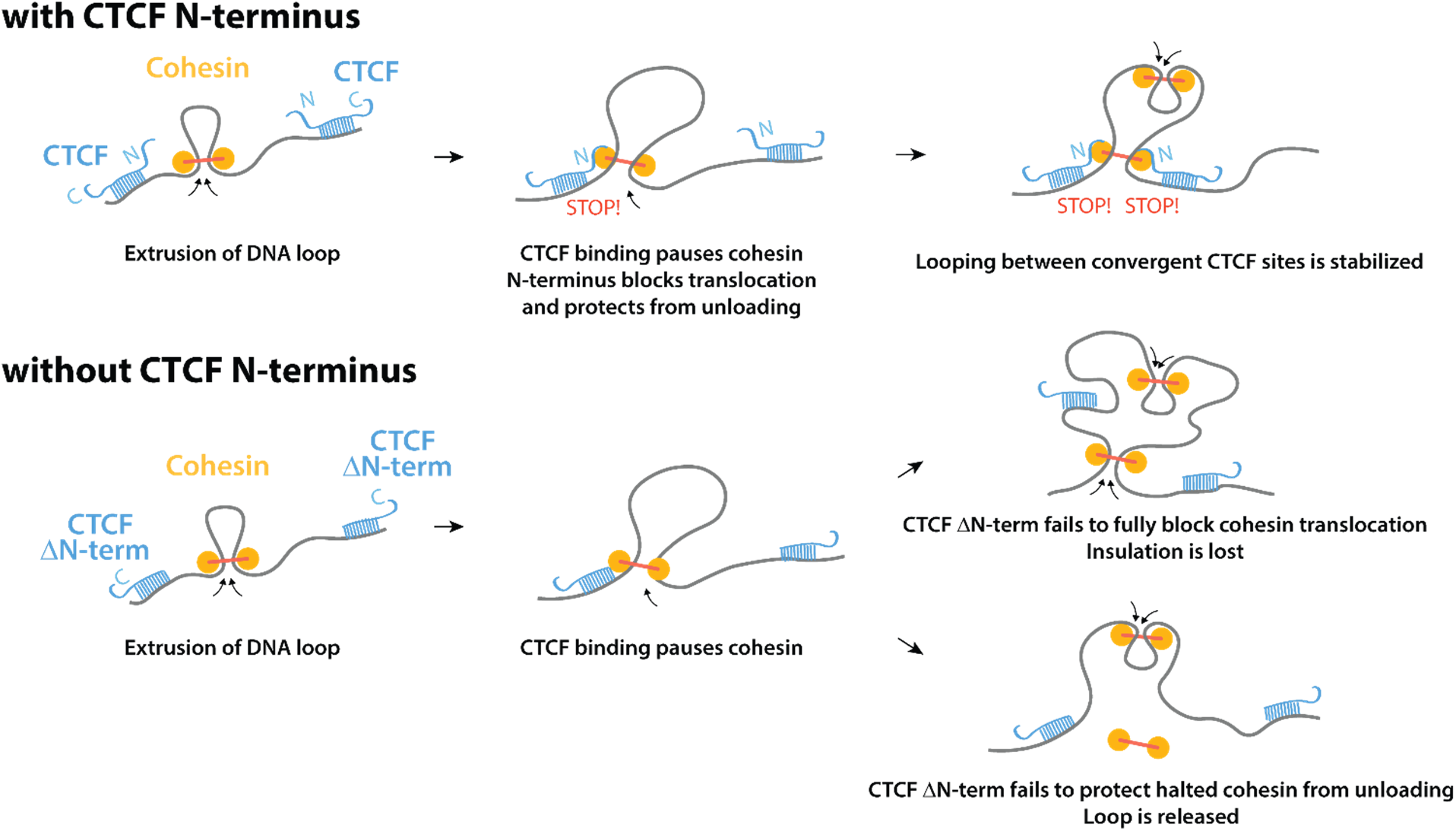
Summary model for the role of the CTCF N-terminus in chromosome folding. Because of the non-palindromic nature of the CTCF DNA motif, the effect of the CTCF N-terminus on cohesin retention and DNA loop stabilization is polarized to one side of CTCF binding site. Altogether, these events results in pairs of interacting TAD boundaries being preferentially populated by CTCF motifs in convergent orientation. Upon deleting the N-terminus of CTCF, cohesin occupancy is diminished but still detectable, indicating that cohesin still pauses upon encountering bound CTCF sites. Loss of cohesin occupancy may reflect either or both decreased ability of truncated CTCF to block cohesin (leading to insulation defects) as well as decreased ability of truncated CTCF to protect halted cohesin from unloading (leading to loss of the DNA loop).

## Supporting information

Supplementary_Table_1_Cell_Lines_and_Vectors

Supplementary_Table_2_sequencing_statistics

---

**Figure 1 – figure supplement 1.**
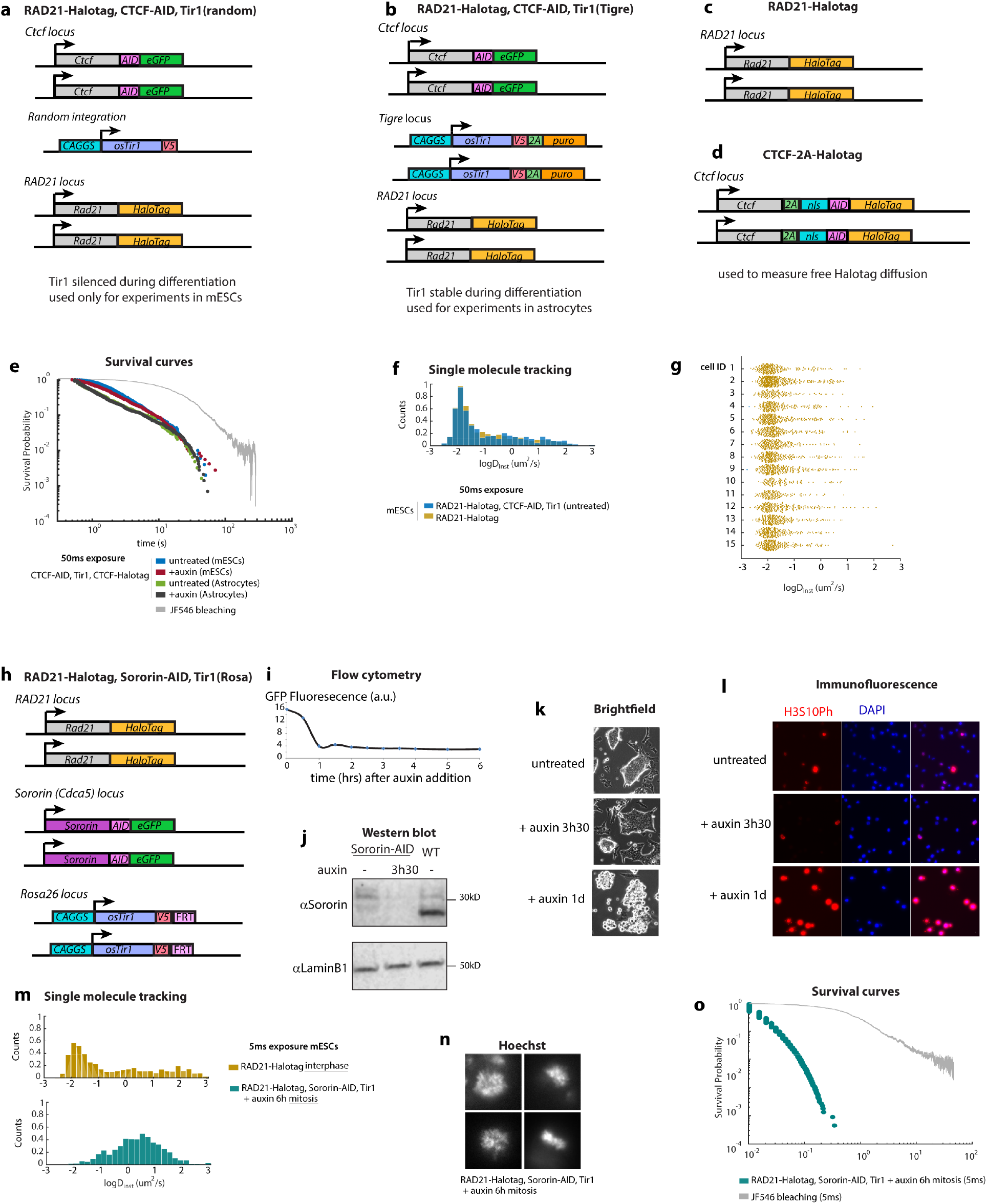
Supporting information regarding RAD21 single molecule tracking in live cells. **a - d**, detailed genotype of the Halotag cells used for live cell single molecule imaging. **e**, survival curves estimated by combining and rescaling data acquired at 5, 50 and 500ms in CTCF-AID, RAD21-Halotag mESCs. Survival Probability computed from data acquired with continuous imaging at 50ms exposure. Statistics: mESC untreated: 507 trajectories; mESC +auxin: 731 trajectories; astrocytes untreated: 628 trajectories; astrocytes +auxin: 1090 trajectories N = 15 cells per condition. **f**, Similar distribution of RAD21-Halotag diffusion coefficient in cells with or without CTCF-AID **g**, RAD21-Halotag single mESCs display similar Diffusion coefficients **h**, detailed genotype of the Sororin-AID-eGFP, Tir1, RAD21-Halotag cells **i**, depletion kinetics of Sororin-AID after auxin addition using flow cytometry for GFP in mESCs **j**, Western blot indicating destabilization of Sororin after addition of the AID tag and complete disappearance after auxin treatment **k**, brightfield imaging of live Sororin-AID-eGFP mESCs illustrating the accumulation of round refringent mitotic cells after incubation with auxin for 1 day (doubling time of parental cells = 12-14hrs) **l**, H3S10 immunofluorescence confirming the accumulation of mitotic cells after auxin treatment of Sororin-AID cells **m**, rapid diffusion coefficients observed mitotic cells **n**, examples of the mitotic figures in cells analyzed **o**, RAD21-Halotag binding events detected in mitotic cells have very short survival time

**Figure 2 – figure supplement 1.**
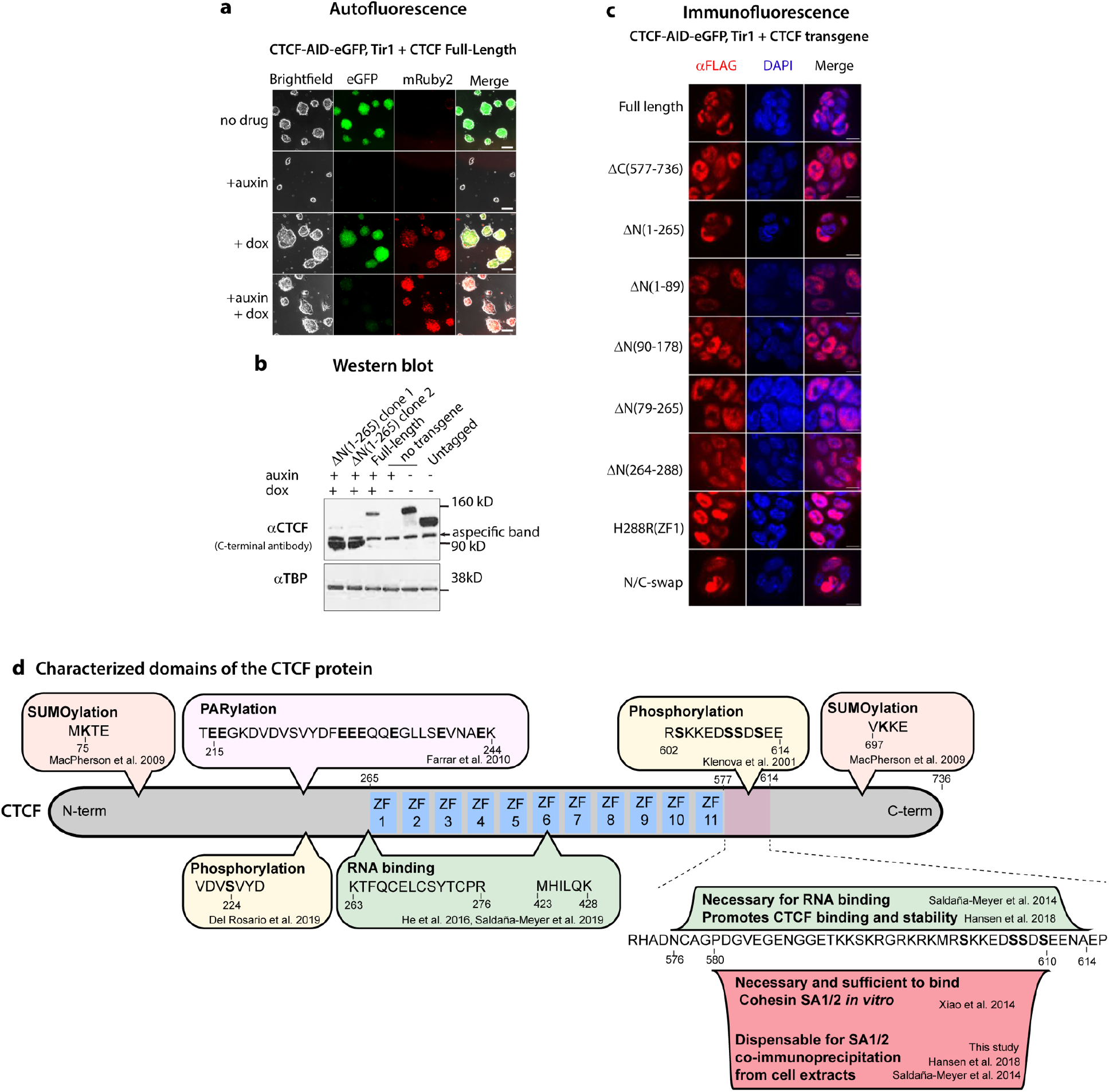
Supporting information regarding the CTCF complementation system. **a**, Brightfield images of live complemented mESCs after 4 days of treatment. scale bar = 100μm. **b**, Western blot with a C-terminal antibody after 2 days of treatment (Milipore 07-729) confirming slightly higher expression of the CTCF Δ(1-265) **c**, all CTCF truncation analyzed displayed nuclear localization using Immunofluorescence against the FLAG tag after 2 days of treatment. scale bar = 10μm. **d**, schematic depiction of known functional and post-translationally modified sites in CTCF. Amino-acid numbers refer to the mouse protein. When information was only available in humans, orthologous amino-acids are reported on the figure.

**Figure 2 – figure supplement 2.**
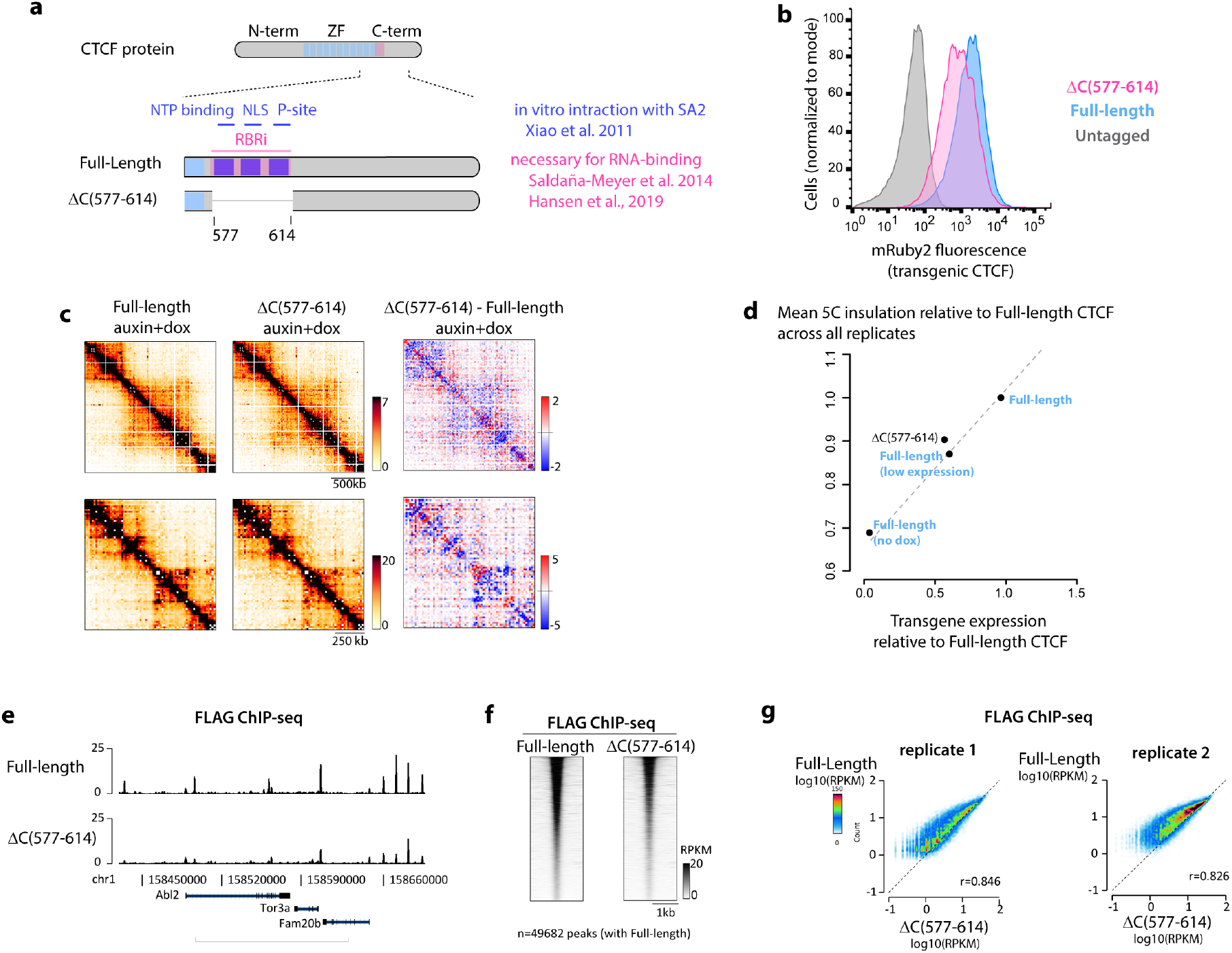
C(577-614) appears dispensable for connecting CTCF and cohesin functionally. **a**, schematic depiction of the position of the Xiao et al. 2011 cohesin SA interaction domain and C-terminal RNA binding region (RBR_i_). **b**, flow cytometry illustrating lower expression level of the ΔC(577-614) transgenic CTCF **c**, 5C snapshots in ΔC(577-614) binned at 15kb. **d**, excerpt of Figure 2b displaying only the ΔC(577-614) mutant. **e-g**, ΔC(577-614) displays lower overall binding to DNA by ChIP-seq than control full-length CTCF with higher transgene expression

**Figure 3 –figure supplement 1.**
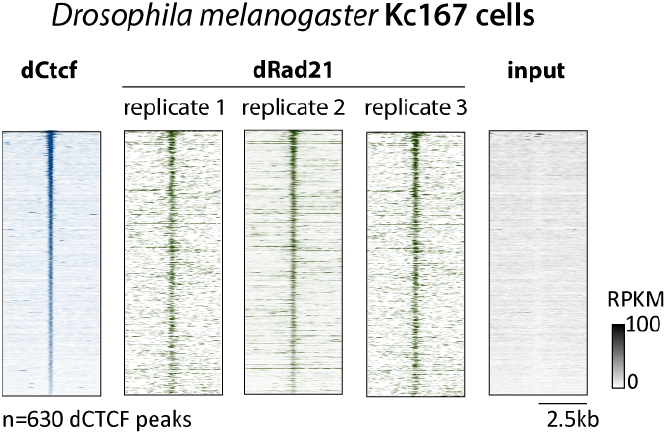
Rad21 is enriched at CTCF binding sites in Drosophila cells. Density plots of CTCF and Rad21 ChIP-seq signal centered at CTCF peaks in Drosophila Kc167 cells(Li et al., 2015).

**Figure 4 - supplement 1.**
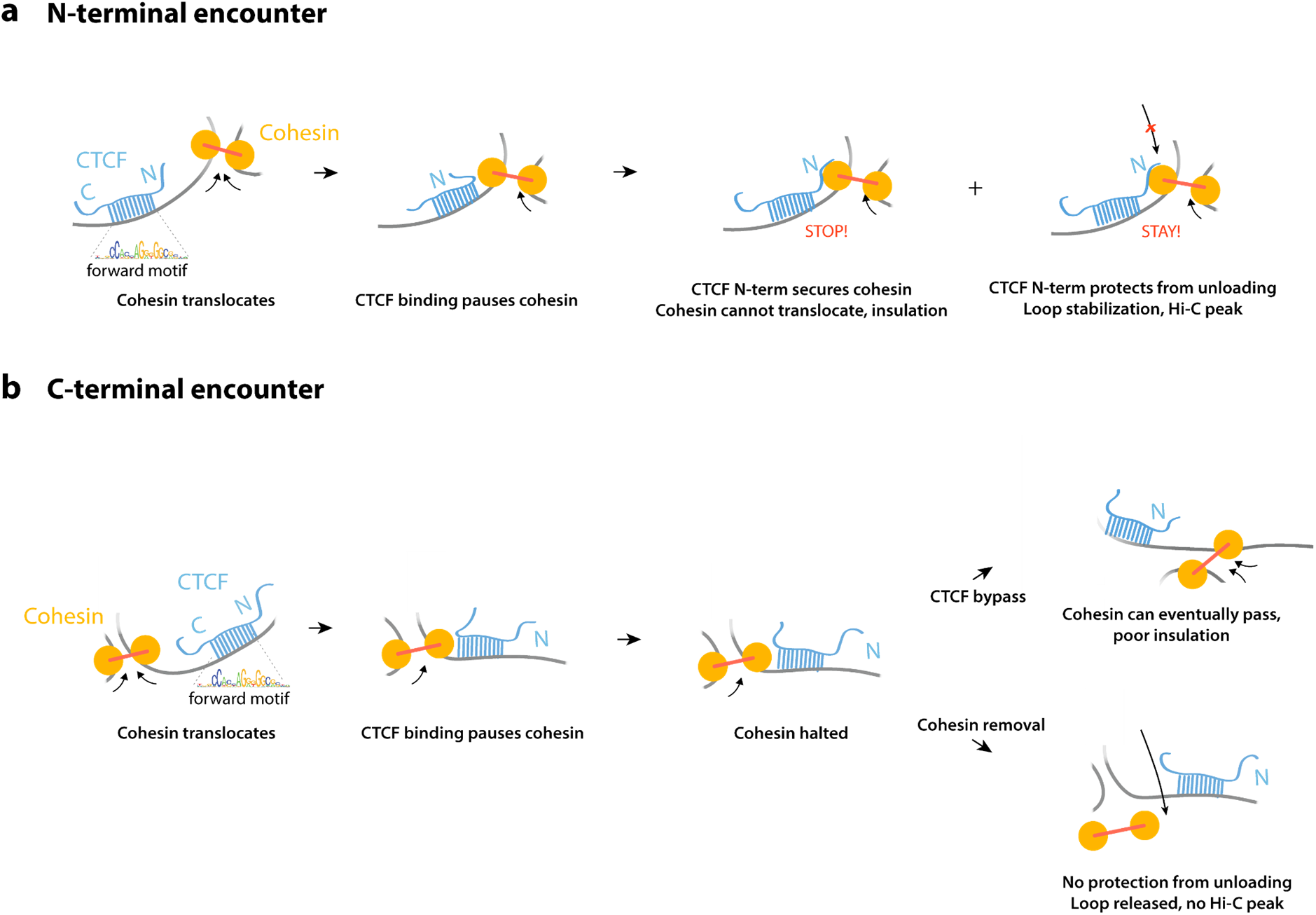
Models for Cohesin encountering either the N- or C-terminal sides of CTCF. **a**, Upon encountering a bound CTCF site cohesin halts, irrespective of motif orientation. Our observations indicate that CTCF then employs multiple amino-terminal subdomains, to block further translocation of cohesin and prevent its unloading. **b**, A similar sequence of events occurs when cohesin encounters normal CTCF from the C-terminal side or N-terminus truncated CTCF from either side: CTCF binding pauses cohesin, and resolution of that pause can involve either eventually bypassing the CTCF site (*e.g*. as CTCF proteins exchange or by passing over), resulting in with no or poor insulation detected by 5C. Alternatively and non-exclusively, paused cohesin is unloaded, likely through PDS5/WAPL, releasing the DNA loop.

## Acknowledgments

This manuscript is dedicated to the memory of Maxime Dahan, who supervised LC. We thank Julia Ronsch, Christel Picard and Edith Heard for support with tissue culture. We are grateful to: Mathieu Coppey for mentoring LC; Luke Lavis, Pierre-Antoine Defossez, Heinrich Leonhardt, David Spector for reagents; Anders Hansen and Maxime Woringer for insight on single-molecule imaging; Mads Lerdrup for support with the Easeq software; Joke van Bemmel and Anton Goloborodko and Erika Anderson for critical reading of the manuscript. We thank the Gladstone Flow Cytometry, Genomics, Histology and Microscopy, and Stem Cell Cores for their excellent services.

## Author Contributions

EPN designed the study with GF, LAM, MD and BGB. EPN created transgenic cell lines, performed and analyzed ChIP-seq, 5C, immunofluorescence, designed and analyzed F3H. LC performed and analyzed single molecule imaging with help of BH, ALS, AC. GF analyzed 5C data and helped in the design of the study. KS, AN, AU provided support for cloning and tissue culture, and KS performed and analyzed F3H as well as Western blots. VK performed ChIP-seq experiments. KSP provided support for bioinformatic analyses. EPN wrote the manuscript with input from all authors, particularly GF. EPN directed the project with BGB.

## Competing interests

none.

**Correspondance and requests for materials** should be addressed to EPN or BGB. Key plasmids will be available through Addgene.

## Methods

### Cell culture

Parental WT mESCs E14Tg2a (karyotype 19, XY; 129/Ola isogenic background) and subclones were cultured in DMEM+Glutamax (ThermoFisher cat 10566-016) supplemented with 15% Fetal Bovine Serum (ThermoFisher SH30071.03), 550μM b-mercaptoethanol (ThermoFisher 21985-023), 1mM Sodium Pyruvate (ThermoFisher 11360-070), 1X non-essential amino-acids (ThermoFisher 11140-50) and 10^4^U of Leukemia inhibitory factor (Millipore ESG1107). Cells were maintained at a density of 0.2-1.5×10^5^ cells / cm^2^ by passaging using TrypLE (12563011) every 24-48h on 0.1% gelatin-coated dishes (Millipore cat ES-006-B) at 37°C and 7% CO2. Medium was changed daily when cells were not passaged. Cells were checked for mycoplasma infection every 3-4 months and tested negative.

To establish neural progenitors and astrocytes, CTCF-AID mESCs were seeded at around 0.1 million cells in a 75cm2 gelatinized dish in mESC medium. The following day cells were rinsed twice in 1X PBS and switched to N2B27 medium (50% DMEM/F12 medium: Gibco 31330-038, 50% Neurobasal medium: Gibco 21103-049, 1X Glutamax Gibco 35050061, 0.5X B27 Gibco 17504-044, 1X N2 Millipore SCM012, 0.1mM 2-mercaptoethanol (ThermoFisher 21985023) and changed daily. After 7 days cells were detached using TryplE and seeded on non-gelatinized bacterial dishes for suspension culture at 3 million cells per 75cm2 and cultured in N2B27 containing 10ng/mL EGF and FGF (Peprotech 315-09 and 100-18B). After 3 days floating aggregates were seeded on gelatinized dishes. After 2-4 days cells were dissociated using Accutase and passaged twice on gelatinized dishes in N2B27+EGF+FGF and cryopreserved after expansion. For differentiation into quiescent astrocytes adherent NPC cultures were washed twice with N2B27 and cultured for at least 48h with N2B27+ 10ng/mL BMP4 (R&D Systems 314-BP-010).

Schneider’s Drosophila Line 2 (S2) cells were obtained from ATCC and cultured in Schneider’s Drosophila Medium (ThermoFisher 21720001) with 10% Heat inactivated FBS (ThermoFisher SH30071.03) at 28°C according to the ThermoFisher protocol.

AID depletion was triggered using 500mM of Indole-3-acetic acid sodium salt (auxin analog) Sigma-Aldrich Cat #I5148 final diluted in culture medium. TetO promoters were induced using of 1mg/ml doxycycline final diluted in culture medium. Single molecule imaging in CTCF-AID cells was performed after 1 day of auxin treatment, to minimize secondary effects. ChIP-seq was performed after 2days of auxin (+dox) treatment to enable comparison with previous ChIP-seq and Hi-C data^2^. 5C was performed after 4 days of auxin (+dox) of treatment, where the effect of CTCF depletion and the difference with the CTCF full-length transgene rescue were maximal^2^.

### Plasmid Construction

Plasmid were assembled using Gibson assemble (SBI MC010B-1) or restriction-ligation. Mouse cDNAs were used for CTCF transgenes, and cloned after by reverse-transcription of mESC mRNAs (SuperscriptIII, ThermoFisher). Targeting vectors driving doxycycline-inducible CTCF cDNAs were assembled by modifying the pEN366 vector^2^.

Parylation-deficient CTCF was created by alanine substitution of the eight glutamic acid residues between position 215 and 244, known to obliterate Parylation^4^. Sumoylation sites were mutated. The N terminal Sumoylation site was obliterated by introducing the previously described^5^ K75R mutation.

The list of plasmids generated in this study can found in the supplementary information. Annotated plasmid sequences are available as supplementary information. Key plasmids will be available through Addgene.

### Genome engineering

For transfection, plasmids were prepared using the Nucleobond Maxi kit (Macherey Nagel) followed by isopropanol precipitation. Constructs were not linearized.

To knock in TetO-CTCF cDNAs at the *Tigre* locus, CTCF-AID, Tir1(random insertion) clone EN52.9.1 (Nora et al. 2017) was transfected using using the Neon system (Thermofisher) using a 100μL tip with 1 million cells at 1400V, 10ms and 3 pulses. 5ug of the *Cas9-Tigre* sgRNA vector pX330-EN1201 (Nora et al. 2017 Addgene #92144) and 15μg of targeting construct. After electroporation cells were seeded in a 9cm^2^ well and left to recover for 48h. Cells were plated at limited dilution and grown for around 7 days in the presence of puromycine at 1μg/mL until single colonies could be picked. Individual clones were genotyped by PCR and analyzed by flow cytometry for induction of the CTCF-mRuby2 transgene on a MACSQuant analyzer. Homozygous clones were identified by PCR and those driving expression as close as possible as the control cells harboring the full-length CTCF transgene were expanded and cryopreserved. See supplementary information.

To knock in the Halotag at RAD21, mESCs (E14Tg2a or CTCF-AID, Tir1(random)) were transfected using using the Neon system (Thermofisher) using a 100μL tip with 1 million cells at 1400V, 10ms and 3 pulses. 5ug of the *Cas9-Tigre* sgRNA vector pX330-EN1082 (see supplementary information) and 15μg of targeting construct pEN313 (see supplementary information). After electroporation cells were seeded in a 9cm2 well and left to recover for 48h, geneticin was then added to the media at 200μg/mL and cells were selected as a heterogenous pool of homozygous and heterozygous cells for around 10 days, at which stage over 70% of the cells showed nuclear fluorescence after addition of fluorescent Halotag ligand. Cells were then transfected with the Neon system using a 10μL tip and 0.1 million cells with 250ng of a flippase-expressing plasmid (pCAGGS-FlpO-IRES-puro)^6^ in order to trigger FRT recombination and excision of the blasticidin selection cassette. After electroporation cells were seeded in a 9cm2 well and left to recover for 48h and transferred into a 78cm2 petri dish from whish two serial 1:10 dilution were seeded in an additional two dishes. After 7-8 days of culture without antibiotic selection single colonies were manually picked, transferred into a 96-well plate, dissociated and re-plated. Clones were then genotyped by PCR for homozygous insertion of the Halotag, checked for geneticin sensitivity, expanded and cryopreserved.

We noticed that the RAD21-Halotag cells derived from the CTCF-AID, Tir1(random), clone EN52.9.1, stopped responding to auxin upon differentiation. We therefore used RAD21-Halotag, introduced Tir1 at the Tigre locus using pX330-EN1201 (addgene #92144) and pEN396 vectors (addgene #92142), and isolated a homozygous knockin clone which we used to introduce an AID-eGFP cassette at both endogenous alleles of CTCF using pEN244 (addgene #92144) and (pCAGGS-FlpO-IRES-puro)^6^. We noticed that when targeted at *Tigre* Tir1 expression remained stable upon differentiation.

To create Sororin-AID cells, RAD21-Halotag cells were were transfected using using the Neon system (Thermofisher) using a 100μL tip with 1 million cells at 1400V, 10ms and 3 pulses. 5ug of the *Cas9-Tigre* sgRNA vector pX330-EN1680 (see supplementary information) and 15μg of targeting construct pEN487 (see supplementary information). A homozygous clone was isolated, used for co-transfection with (pCAGGS-FlpO-IRES-puro)^6^ to remove the blasticidin selection cassette. Tir1 was then introduced at rosa26 using vectors pX330-EN479 (addgene #86234) and pEN114 (addgene # 92143). Homozygous clones were identified by PCR.

The list of cell lines generated in this study and the corresponding CRISPR sgRNAs can found in the supplementary information.

### Live single molecule imaging

#### Microscopy set up

Single molecule imaging was performed on an epifluorescence inverted microscope (IX71, Olympus) in HILO illumination^7^. 500mm achromatic lens conjugates the slit to the specimen plane to achieve a proper HILO. The lens focuses the excitation beam on the back focal plane of a 150X objective lens (UApo N 150X TIRF 1.45 NA, O.I., Olympus, France). The lens is mounted on a translation stage together with a metallic mirror that sends the beam to the microscope. Displacement of the translation stage allows a precise positioning of the focused beam at the back focal plane of the objective without influencing the lens-BFP distance. Thanks to this configuration it is possible to adjust the tilting of the laser beam at the output of the objective and thus the effective thickness of the tilted light sheet excitation at the specimen.

Efficient separation between the excitation and emission was achieved with a fluorescence cube containing a quad band dichroic mirror (FF409/493/573/652-Di02-25×36, SEMROCK) together with adequate emission filters. The setup is provided with a 561 nm laser (Sapphire 561, Coherent, Santa Clara, CA, USA), a 488 nm laser (488LM-200, ERROL, France) and a 405 nm laser (405LM-200, ERROL, France). Lasers were tuned via an acousto-optical tunable filter (AOTFnC-400-650-TN, A&A Optoelectronic, France) and controlled by a home-made interface in Micromanager (Edelstein et al, 2014). Signal was acquired with an EM-CCD camera (iXonEM DV860DCS-BV, Andor, Ireland) run in frame-transfer mode

#### Acquisitions

To perform single molecule tracking experiments, cells (both mESC and Astrocytes) were grown on circular petri dishes with glass bottom (MatTek, Part No: P35G-1.5-14-C) coated with fibronectin (Millipore SAS cat# FC010-5mg). Cells were seeded at a density of 3×10^5^/cm^2^ the day before the experiments, in culture medium based on Fluorobrite DMEM for mESCs (ThermoFisher A1896701) and in phenol-red N2B27 with BMP4 for astrocytes (ThermoFisher 12348017). We underline the importance of performing single molecule imaging in phenol-red free medium to both reduce the background fluorescence and minimize localization errors.

The experiments were performed 20 hours (labeled as 1 day) after adding auxin to culture medium. To achieve single molecule labelling cells were incubated with 1pM of Halo-JF549 for 20 minutes at room temperature (incubation followed by a first rinsing step, 15 minutes wait and another rinsing). While waiting for the second rinsing step cells were incubated with 1□M Hoechst and consequently washed to minimize the fluorophores unbound in solution. All washings were performed using cell culture medium; the coverslips treated with auxin were washed with medium enriched with auxin. During the experiments cells were kept at 37° and 5% CO2 with a Tokai Hit heating system (INUBG2E-PPZI).

To locate nuclei, cells were stained with Hoechst 33342 (bisBenzimide H 33342 trihydrochloride, Sigma-Aldrich, ref 14533), excited with 405 nm light. The CTCF-GFP was imaged in the 488 nm channel. To track Cohesin-Halo-JF549, the sample was excited with the 561 nm laser. At least 5000 frames were recorded in a continuous imaging regime, the laser being controlled by the camera. Laser power was approximately 0.1 kW/cm^2^ and adjusted depending on the exposure time in order to keep the amount of excitation photons constant.

To determine the fraction of bound molecules we acquired images in a continuous regime at a frame rate of 197Hz (5ms). For the analysis of the dynamics (MSD) and the residence time we acquired videos at a rate of 20Hz (50ms).

#### Quantification of photobleaching

To characterize the photobleaching of the organic dye used for our Single Particle Tracking experiments (SPT), we acquired movies in the same imaging conditions of the SPT experiments in terms of laser power and exposure. Cells were stained with the JF549 organic dye^8^ at 1nM for a bulk labelling. The plot in **Figure 1 – figure supplement 1** shows the average normalized bleaching curve for acquisitions made with an exposure time of 50ms with the same laser power used for the SPT experiments.

### Analysis of single particle tracking data

To localize the single emitters and build the trajectories we used SLIMfast^9^, implemented in Matlab and based on the MTT algorithm^10^. The Point Spread Function (PSF) of a single emitter is fitted with a 2D-gaussian, whose center corresponds to the position of the fluorophore with a sub-pixel resolution.

#### Analysis of bound fractions

To quantify the fraction of bound molecules we used data acquired at 5 ms exposure in a continuous imaging regime. The actual framerate acquisition is 197Hz (5.08 ms), due to the frame transfer lag to the camera. We chose to use the data from the fastest acquisition rate to include the fastest diffusing population, which blurred when imaging with 50ms exposure time.

Particles were tracked as described above and we computed the distribution of the step sizes of the protein of interest. The trajectories consisted of at least one step, or 2 localizations. A two-state model was chosen to fit our data. The computation of the fraction of bound molecules is corrected for the subset of free molecules that may leave the focal plane^11^. The fit was performed on the Cumulative Distribution Function (CDF) to avoid biases due to the binning choice.

#### Residence times

To further characterize the binding kinetics, we extrapolated the trajectories that stayed confined in a circular area of radius r = 2 pixels for the whole duration. With this pool of “immobile” trajectories we built the distribution of residence times and consequently computed the Survival Probability. Such distribution of residence times is defined as the inverse cumulative probability, or the probability for a molecule to have a life longer than 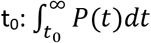

Given the intrinsic limitations of single molecule imaging when probing very stable binding events (as for Cohesin) we use the Survival Probability curves to qualitatively sample the discrepancies between the different biological conditions.

#### Analysis of diffusion dynamics

The trajectories obtained from experiments at 50ms were analyzed with custom codes implemented in Matlab. First, we computed the time-averaged Mean Squared Displacement (MSD) as MSD=<xt+nΔt-xt>, where x(t) is the position at time point t; n = 1, 2 …, N, with N = maximum number of time points in a trajectory, and <> indicating the ensemble average over all the possible time lags of one individual trajectory.

We selected the trajectories with at least 10 localizations. In spite of the low JF549 ligand concentration the beginning of the videos are very dense in point emitters. We therefore cut the first hundred frames of the raw movies, and we only performed tracking on images with approximately 10 molecules per frame. We did not threshold data used to quantify the fraction of bound molecules nor to the estimation of the Survival Probability.

Once computed the MSD we extrapolated what we call the instantaneous diffusion coefficient (D_inst_) from each trajectory by fitting the MSD from point 2 to point 6. We followed the common approach of performing a linear fit, assuming a purely Brownian motion at the beginning of the MSD^9,12^.

#### Detailed statistics

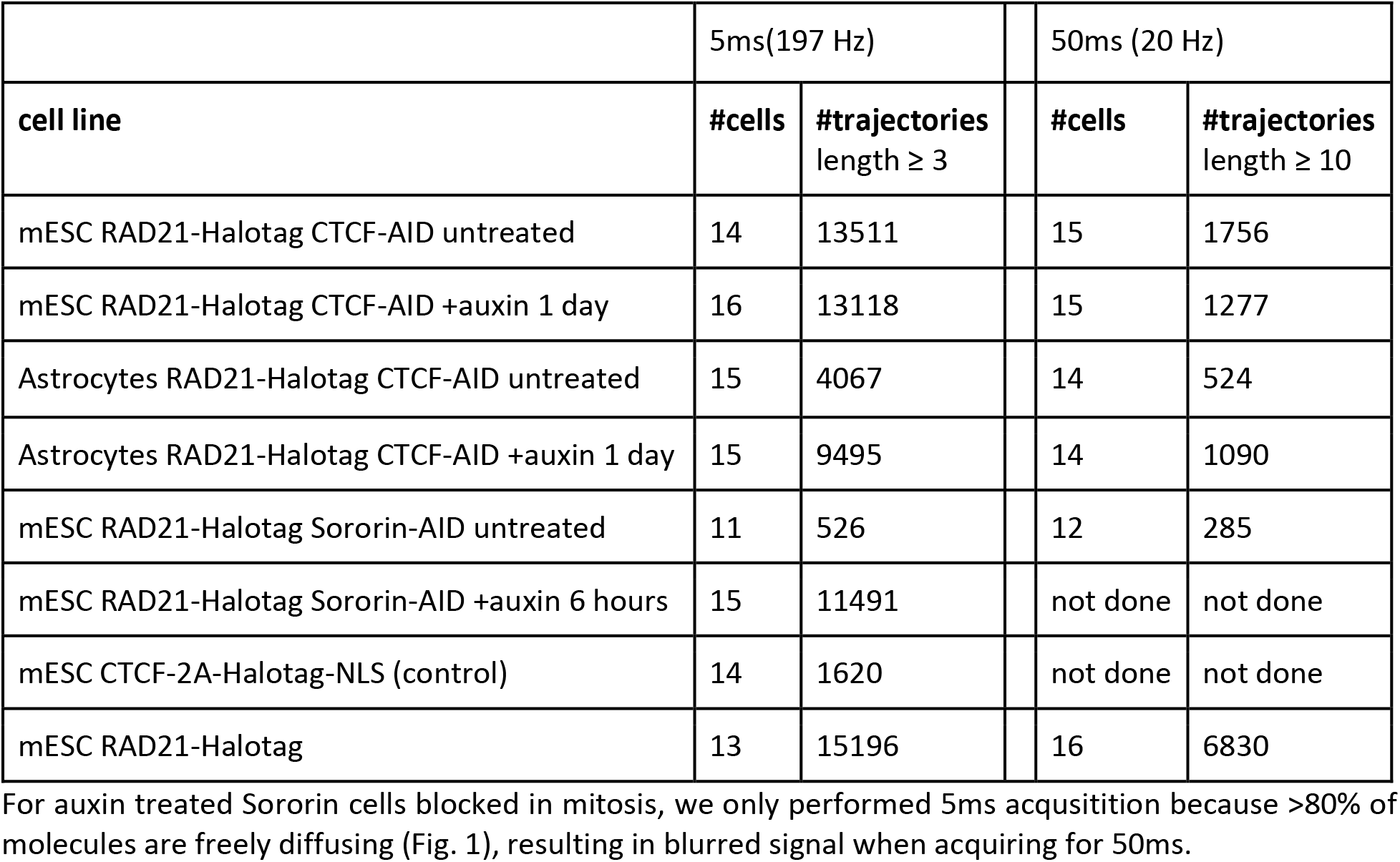

Statistics related to **Figure 1 – figure supplement 1e**

| Cell ID | #trajectories |
| --- | --- |
| 1 | 165 |
| 2 | 157 |
| 3 | 62 |
| 4 | 150 |
| 5 | 118 |
| 6 | 94 |
| 7 | 76 |
| 8 | 141 |
| 9 | 61 |
| 10 | 107 |
| 11 | 115 |
| 12 | 63 |
| 13 | 126 |
| 14 | 77 |
| 15 | 130 |

### Immunostaining

mESCs were grown on glass-coverslips, fixed with 3% formaldehyde in 1XPBS for 10’ at room temperature. Permeabilization was carried out in 0.5% Triton followed by blocking with 1% BSA diluted in 1X PBS (Gemini cat 700-110) for 15min at room temperature. Primary antibody incubation was performed at room temperature for 45min (Monoclonal ANTI-FLAG^®^ M2 antibody produced in mouse Millipore-Sigma F1804 at 1/250 dilution), followed by three 5 min washes in 1X PBS, secondary antibody incubation (AlexaFluor594 Goat anti-Mouse IgG Invitrogen A-11005 at 1/10.000 dilution), three 5 min washes in 1X PBS, counter-staining with DAPI and mounting in 90% glycerol – 0.1X PB – 0.1% p-phenylenediamine pH9. Images were acquired on a Zeiss spinning disk with 60X objective. In order to avoid loss of loosely-attaching mitotic cells for the H3S10 immunostaining in Sororin-AID cells, cells were detached with TryplE, spun in culture medium, resuspended in PBS and let to attach for 10min in 1XPBS 25μl droplets spotted onto 0.1% poly-L-Lysine coated coverslips. Cells were then processed as described above, except that the primary antibody used was Anti-H3S10Ph, rabbit polyclonal Millipore 05-636.

### Flow cytometry

mESCs were dissociated with TryplE, resuspended in culture medium, spun, and resuspended in 4% FBS-PBS before live flow cytometry on a MACSQuant instrument (Miltenyibiotec). Dissociation, wash, and flow buffers were supplemented with auxin, when appropriate, to avoid re-expression of the CTCF-AID-eGFP fusion. Analysis was performed using the Flowjo sowftware.

### Western blots

mESCs were dissociated, resuspended in culture medium, pelleted, washed in PBS, pelleted again and kept at −80°C. 15-20 million cells were used to prepare nuclear extracts. Cell pellets were resuspended in 10mM HEPES pH 7.9, 2.5mM MgCl2, 0.25M sucrose, 0.1% NP40, 1mM DTT, 1X HALT protease inhibitors (ThermoFisher) and swell for 10 min on ice. After centrifugation at 500 g nuclei were resuspended in on ice in (25mM HEPES pH 7.9, 1.5mM MgCl2, 700 mM NaCl, 0.5mM DTT, 0.1 mM EDTA, 20% glycerol, 1mM DTT, 250 U Benzonase and incubated on ice for 10min. Insoluble materials were pelleted by centrifugation at 18,000 g at 4°C for 10 min and supernatant (nuclear extracts) were stored at −80C. Protein concentration from supernatants were measured using the Pierce Coomassie Plus assay kit (Thermofisher).

For CTCF Western blot in **Figure 2 – figure supplement 1**, 40ug of nuclear extracts were loaded per lane. Samples were mixed with Laemmli buffer and 2.5% beta-mercaptoethanol, then loaded onto a Bolt 4%-12% Bis-Tris Plus gel (ThermoFisher). Gels were wet transferred onto PVDF membranes in transfer buffer (25mM Tris-Base, 192mM Glycine, & 10% Methanol) for 3 hours at 80V. Membranes were blocked for 2 hours with Odyssey blocking buffer (Li-Cor cat. 927-40000) and subsequently incubated with primary antibody overnight at 4°C (1:1000 anti-CTCF C-terminus Millipore 61311 and 1:2000 anti-TBP Abcam ab51841) in Odyssey blocking buffer. Membranes were washed three times in TBT-0.1% Tween, 5-10 minutes per wash, and were incubated with secondaries at room temperature for 1 hour (1:10000 HRP-anti-rabbit Cell Sig #7074 and 1:10000 HRP-anti-mouse Cell Sig #7076). Blots were wash 3 times for 5-10 minutes in TBS-0.1% Tween. CTCF blot used Amersham ECL Prime Western Blotting Detection Reagent (GE RPN2236) and TBP blot used Amersham ECL Western Blotting Detection Kit (GE RPN2108) for HRP activation. Blots were then exposed onto x-ray films for different exposure times.

### ChIP-seq

#### Preparation of Spike-in chromatin from S2 cells

Cells were detached from culture dished by splashing them gently but thoroughly with culture medium, and transfered to 15mL conical tube **before** spinning 1000g for 3 min. Cells were resuspended at 10^6 cells/mL in complete S2 culture medium at room temperature. 270uL of 37% Formaldehyde (Electron Microscopy Sciences) for a final concentration of 1%, and agitated on an orbital shaker for 10min @ RT. 510uL 2.5M glycine (final concentration 125mM) was added and cells were lef agitating for 5min @ RT, then spun at 1000g for 2min, 4C. Fixed cells were wash once in 1mL cold 1XPBS-0.125M Glycine, Spin at 1000g for 3min, 4C. Cells were used for sonication without prior freezing, as we noticed that snap freezing dramatically altered shearing efficiency. Fresh cell pellets containing were resuspended in 1mL Cell lysis buffer (20mM Tris HCl pH8.0, 85mM KCl, 0.5% IGEPAL and 1X Halt protase inhibitors ThermoFisher PI78425),) and incubated on ice for 10min. Nuclei were pelleted by spinning at 2500g for 5min at 4C and lysed in 50mM Tris HCl pH8.0, 10mM EDTA, 1% SDS and 1X Halt protase inhibitors for 30min on ice. Chromatin was sheared using a Covaris S220 ultrasonicator 5% Duty cycle, 5 intensity and 200 cycles/burst for 7min. Debris were pelleted by centrigugation at 1500g for 5min. Supernatent was transferred in a new tube and glycerol was added to 10% final before freezing at −80C as single use aliquots. For each ChIP experiment, 600ng of Drosophila chromatin (as estimated from the amount of DNA retrieved after reverse-crosslinking) was used in combination with sonicated chromatin obtained from 10 million mESCs.

#### Rad21 ChIP-seq in Figure 1

The first set of Rad21 ChIP-seq was performed in parallel of CTCF ChIP-seq in the CTCF-AID mESCs clone EN52.9.1 published in 2017^2^, using 10mg of antibody abcam ab992 together with 40ng of Drosophila melanogaster spike-in chromatin (Active motif 53083) and spike-in antibody (Active motif 61686). These tracks are tagged as “2017protocol” in the supplementary table and companion GEO submission of this study.

#### Rad21 and FLAG ChIP-seq in Figure 3

FLAG and Rad21 ChIP-seq in mESCs containing CTCF rescue transgenes, as well as replicates of the parental CTCF-AID line EN52.9.1 post 2017, were prepared with a protocol differing from data in Figure 1 by the lysis and wash buffers. For the full-length transgene we used the high-expressing clone (EN133.10) to be closest to the expression level of the ΔN(1-265) clones.

For fixation, mESCs were dissociated using TrypLE and resuspended in 10% FBS in PBS, counted and adjusted to 1 million cells per mL. Formaldehyde was then added to 1% final followed by 10 min incubation at room temperature. Quenching was performed by adding 2.5M Glycine-PBS to 0.125M final followed by 5 min incubation at room temperature, 15 min incubation at 4°C, centrifugation at 200 g 5 min at 4°C, resuspended with 0.125M Glycine in PBS at 10 million cells per mL, aliquoted, spun at at 200 g 5 min at 4°C and snap frozen on dry ice.

Fixed cells were thawed on ice, resuspended in ice cold 20mM Tris HCl PH8.0, 85mM Kcl, 0.5% IGEPAL and 1X HALT protease inhibitor, counted and readjusted to obtain 10 million cells total exactly, incubated on ice 15 min, centrifuged at 500 g 5 min at 4°c, resuspended in 1mL 20mM Tris HCl pH8.0, 0.1% SDS, 0.5% Sodium Deoxycholate and 1X HALT protease inhibitor, transferred to a MilliTube (Covaris). Chromatin was sheared on a Covaris S2 sonicator for 15 cycles at 5% duty cycle, intensity 8, 200 cycles per burst in a waterbath maintained at 4°C, using 1 min sonication – 30 s rest, resulting in fragments. Samples were clarified by centrifugation at 18,000 g at 4°C for 10 min. Supernatents were transferred to 15mL conicals and 600ng of spike-in Drosophila chromatin (home made) was added. 10% of the mixture was saved as input and the rest was diluted to 5mL with ice-cold 16.7mM Tris-Hcl pH 7.4, 167mM NacCl, 0.01% SDS, 1.1% Triton X-100, 1.2mM EDTA, 1X protease inhibitor. 10μg of anti-FLAG (Millipore-Sigma F1804) or anti-Rad21 (Abcam 992) together with 4μg spike-in antibody (anti-H2Av, Active motif) together with 4μg spike-in antibody (Active motif) was added alongside with 40μL prewashed protein G Dynabeads (ThermoFisher) followed by overnight incubation at 4°C on a rotator. Beads were collected using a magnetic stand, transferred into 2mL tubes and washed with 1mL twice 5min with 20mM Tris HCl pH 8.0, 150mM NaCl, 2mM EDTA, 0.1% SDS, 1% Triton X 100, twice 5min with 20mM Tris HCl pH 8.0, 500mM NaCl, 2mM EDTA, 0.1% SDS, 1% Triton X 100, twice 5min with 10mM Tris HCl pH 8.0, 0.25M LiCl, 1mM EDTA, 1% NP40, 1% Sodium Deoxycholate, and rinsed twice with 1X TE buffer. DNA was eluted twice by resuspending washed beads with 50uL 1% SDS, 0.1M NaHCO3 and incubating for 30min and pooling eluates. Saved input DNA was diluted in the same buffer and treated similarly. 1mL of 10mg/ml DNAse free RNAse A was added and eluates were incubated at 37C for 30 minutes, prior to addition of Add 1ul of 20mg/ml Proteinase K and 12ul of 5M NaCl and overnight incubation at 65C. The next day DNA was clean either using Ampure Beads (FLAG ChIPs) or Qiagen PCR cleanup minelute kit, eluting in 32mL. DNA was then used for library preparation exactly as described^2^, using the entire eluate for ChIP-seq and 40ng for inputs.

### ChIP-seq analysis

Mapping and peak calling was performed as exactly as described previously^2^. Published^2^ CTCF ChIP-seq peaks in untreated and auxin-treated CTCF-AID mESCs were used to identify total and auxin-sensitive CTCF peaks. Fraction of reads in peaks (FRIP) scores were calculated by calculating the proportion of uniquely mapping reads within auxin-sensitive CTCF peaks compared to the total number of uniquely mapping reads, and excluding genomic regions known to display artificial ChIP-seq signal^13^ retrived from https://sites.google.com/site/anshulkundaje/projects/blacklists

Published^14^ datasets from accession GSE63518 were mapped to dm3 and peak calling was performed as exactly as described previously^2^. Data presented in figure 1 were obtained using the commercial Active motif spike in reagents where spike-in calibration yielded consistent results. For Rad21 ChIP-seq in mESCs with the CTCF transgenes (figure 3) we noticed that spike-in normalization gave inconsistent results, artificially rescaling up or down Rad21 scores beyond reason and inconsistently between replicates. To avoid these artifacts we display FLAG and Rad21 analyses without recalibration. Reads were mapped separately to mm9 and dm3 as described^2^, eliminating low-quality reads, PCR duplicates and multimapping reads. Tracks and density plots were generated using Easeq^15^ http://easeq.net/.

### Chromosome Conformation Capture Carbon-Copy (5C)

5C was performed exactly as described^2^.

### 5C analysis

Sequencing and mapping was performed as described^2^. Matrices were then iteratively corrected at the fragment level and normalized to sum to 1e6. Iterative Correction was performed on raw unbinned matrices (fragment level from the alternating 5C primer design) using iterative_correction_asymmteric with default values, (cooltools, https://github.com/mirnylab/cooltools). 5C heatmaps data depicted in the figures were obtained after binning the corrected matrices at 15kb by taking the median over all primer pairs that fall within each pair of bins.

To minimize possible artifacts when calculating insulation scores, we binned the matrices at 20kb by taking the mean over all primer pairs that fall within each pair of bins. The first two diagonals of the binned matrix were then filled with the mean of the second diagonal. Combined insulation scores for each sample were calculated for the binned corrected matrices by aggregating over the same set of boundary positions across samples. Boundaries were identified in untreated CTCF-AID mESCs without any CTCF transgene (GEO accession GSE98671 samples GSM2609248, GSM2609253 and GSM2609256)^2^ by taking the minima of the insulation profile, as previously^2^. Insulation scores were calculated with a 100kb window, as previously^2^. These minima were then filtered to exclude those that are shared with those upon auxin-mediated degradation of CTCF-AID for 4 days in mESCs (GSM2609254, GSM2609259) (to eliminate CTCF-independent boundaries - *e.g*. compartment transitions). Combined insulation scores averaged across all replicates (Fig. 2) were calculated as the mean across boundary positions and averaged across replicas, for each cell line separately. To calculate insulation relative to Full-length transgenes, averages of mutant cDNAs were divided by the average obtained with the reference Full-length transgene. The genomic position of the CTCF-dependent boundaries used were:

| chr | start | stop |
| --- | --- | --- |
| chrX | 99151148 | 99171148 |
| chrX | 99411148 | 99431148 |
| chrX | 100451148 | 100471148 |
| chrX | 100671148 | 100691148 |
| chrX | 101211148 | 101231148 |
| chrX | 103211148 | 103231148 |

Similar results were obtained when using the four most visually prominent boundaries.

Differential heatmaps were generated by binning each matrix independently and subtracting the 5C counts from the reference matrix.

